# Asymmetry in hydrophobicity induces electric potential in non-charged protein condensates

**DOI:** 10.1101/2025.10.15.682625

**Authors:** Leshan Yang, Wen Yu, Xiangze Zeng, Yifan Dai

## Abstract

The capacity of biomolecular condensates to establish and modulate electrochemical equilibria is emerging as an important functioning mechanism in cellular biochemistry. However, the physical chemistry basis of the electric potentials arising from biomacromolecular phase transitions remains unclear. Here, we show that asymmetry in hydrophobicity, which is a generalizable feature in condensate system, can directly encode an electric potential gradient between the dilute and the dense phases. We demonstrate that using a non-charged intrinsically disordered protein, ion-dependent kosmotropic effect can encode measurable pH and interphase potential gradients into condensate. All-atom molecular dynamics simulations further reveal that the distinct intrinsic transfer free energy of ions defines the ion partitioning capability of condensates via favorable interactions with protein backbones. The simulation also shows the existence of both interfacial and interphase electric potentials. These built-in potentials modulate the partitioning and reactivity of charged solutes, enabling non-enzymatic, potential-dependent chemistry within condensates. Our findings identify hydrophobic asymmetry as a simple and generalizable mechanism for charging biological matter, linking water activity and ion energetics to the emergent electrochemistry of condensates.

## Introduction

Condensed biological matter, such as biomolecular condensates, arises from the phase transitions of biomacromolecules, regulating diverse cellular processes, from stress responses to organizing neuronal signaling^1–4^. An emerging feature of such condensed biological matters is their inherent electrochemical activity^5–8^, which is largely driven by the electric potential gradients between the dilute and the dense phases established by the phase transition process^6,8–10^. Recent works have shown that such inherent electrochemical activity of condensates can drive interfacial electric field-dependent redox reactions^6,8,11–13^. Such non-enzymatic, spontaneous redox activity of condensates has been demonstrated to be critical to drive the aggregation of neurotoxic proteins^7,14^, thereby directly contributing to the developments of neurodegenerative disorders. Though such electrochemical functions of condensed matter are generalizable and important to cellular functions^11,12,15–19^, our understanding of the mechanism on how the electrical potential is established through biomacromolecular phase transition is limited.

Polymer theories have shown that the establishment of electric potential gradient between phases can be induced by asymmetry of certain physical quantities in phase transition^20–22^. For example, the concentration asymmetry of polycation and polyanion defines the charge asymmetry in the dense phase^21^. Enforced by charge neutrality within a phase, such asymmetry drives the partitioning or exclusion of small ions into or from the phase^5,10^. Under electrochemical equilibrium between the two phases, the asymmetric distributions of charged species collectively define the Galvani potential^20,21^. Thus, it is commonly assumed that such asymmetry should be initiated by charged polymers, like polyelectrolytes^23–25^, which mediate phase transition through associative or segregative transition with solvent and solute molecules^26^. However, a critical physical chemistry property that is overlooked in driving such asymmetry is water environment itself.

In this study, we explored whether electric potential gradient can be induced by non-charged polymers through encoding asymmetry in hydrophobicity between the dilute and the dense phases. To this end, we implemented phase-separable intrinsically disordered proteins containing only uncharged residues. Our hypothesis is based on the following rationale. Phase transition of IDPs, in which solvent-IDP interactions switch to IDP-IDP interactions, can mediate water concentration gradient and water activity difference between phases^11,19^. Such differences in water concentration and water property encode a unique dense phase environment, such as micropolarity^17,27,28^, which essentially is a manifestation of the hydrophobic solvent environment in the condensates. We therefore asked whether the asymmetric hydrophobicity between the dilute and the dense phases could influence ion distributions during phase transition and whether the ion gradient is substantial enough to encode an electric potential gradient that could modulate condensate electrochemical activity. We reasoned that because different types of ions have distinct hydration free energies, they would respond differently to the distinct hydrophobic environments of the dilute and the dense phases, leading to differential partitioning and the formation of an ion concentration gradient, which can possibly establish an electric potential gradient^29,30^.

Here, we demonstrate the asymmetric hydrophobicity-dependent charging effect using elastin like polypeptide (ELP), which does not possess any charged residues and undergoes phase separation driven by hydrophobic interactions^31–38^. We discovered that the phase transition of ELP can encode an ion concentration gradient depending on the types of ions in the solution system. We investigated the roles of non-specific ion adsorption on the ELP, dense phase water environment and ion transfer free energy on defining the interphase electric potentials of ELP condensates. We show that asymmetry in hydrophobicity serves as an orthogonal driving force to charge biomolecular condensates, transforming non-charged condensates into electrochemically active condensates, which can power inherent electrochemical functions. For the same types of inherent chemical functions, charge condition of condensates can mediate chemical selectivity. Our study uncovered that non-charged IDPs exploit hydrophobicity and kosmotropic effects of ions to encode interphase electric potentials of biomolecular condensates, delivering a new fundamental mechanism understanding the electrochemistry and inherent chemical activities of condensates.

## Results

### Salting out ELP mediates a pH gradient between phases based on chaotropic effect

We used the ELP^39^, [Val-Pro-Gly-Val-Gly]_60_, as the model IDP for this study. The phase transition of ELP is driven by hydrophobic interactions and can be triggered by a salting out process (**Figure 1A**), during which increasing solution salt concentration drives condensate formation. To evaluate whether electric potential could be established, we set out to measure the pH gradient between phases (**Figure 1B**). As hydroxyl ions and hydronium ions reflect the Galvani potential under the electrochemical potential equilibria between phases^8,30^, the pH gradient between phases represents the difference of the electric potentials between phases. To measure pH gradient, we applied C-SNARF-4, which is a ratiometric pH probe allowing us to quantify apparent pH between phases in a dye-concentration independent manner. The application of this probe has been well-calibrated for the use in biomolecular condensates^6,40,41^. We observed a pH gradient between the dilute and dense phases at a salt concentration of NaCl at 500 mM (**Figure 1C**); however, the gradient is limited compared to those established by charged IDPs^6,41,42^. This suggests that the ELP itself might not contribute to environmental pH through charge regulation effect by the side chain^43^, implying its limited contribution to the interphase electric potential.

**Figure 1.**
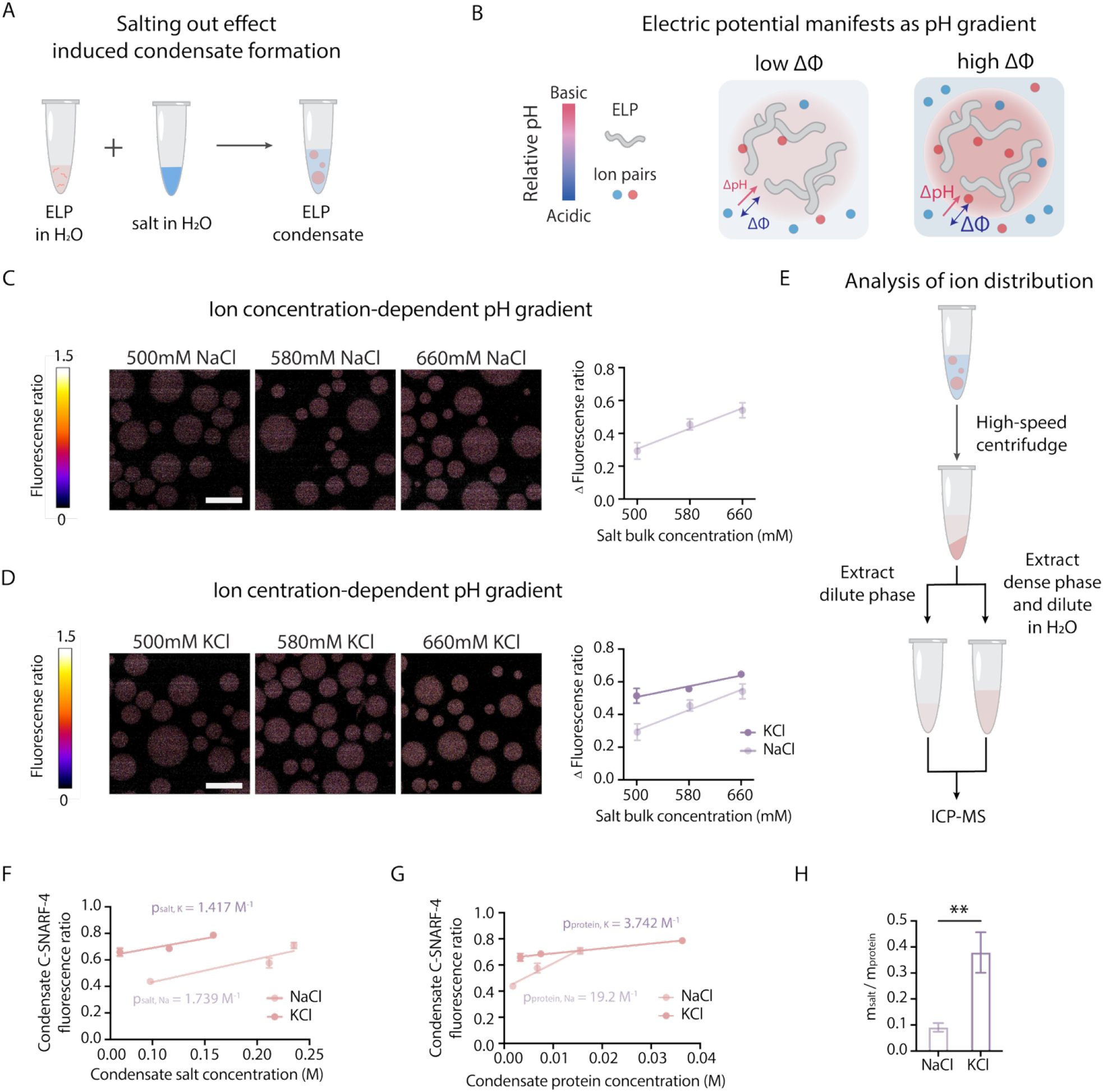
Interphase pH gradient induced by ELP phase separation is modulated by kosmotropic ion effect. **(A)** Schematic of ELP condensate formation. A concentrated salt solution is added to an aqueous solution of ELP protein, triggering phase separation and condensate formation at room temperature. **(B)** Conceptual model illustrating how ion partitioning regulates the interfacial pH gradient. Increasing the bulk salt concentration enhances the ion concentration gradient between the dense and dilute phases, leading to a steeper pH gradient. **(C)** Representative C-SNARF-4 ratiometric fluorescence images of ELP condensates es formed at 500, 580, and 660 mM of NaCl. The rainbow color scale (blue to red) indicates acidic to basic pH conditions. Δ Fluorescence ratio denotes the difference in fluorescence ratio between the dense and dilute phase. Data represent mean ± s.d. from n = 3 independent samples; >10 condensates of similar size were analyzed per condition. Scale bar, 20 µm. **(D)** Representative C-SNARF-4 ratiometric fluorescence images of ELP condensates es formed at 500, 580, and 660 mM of NaCl and KCl. The rainbow color scale (blue to red) indicates acidic to basic pH conditions. Δ Fluorescence ratio denotes the difference in fluorescence ratio between the dense and dilute phase. Data represent mean ± s.d. from n = 3 independent samples; >10 condensates of similar size were analyzed per condition. Scale bar, 20 µm. **(E)** Schematic of separating the dense and dilute phase of solution containing ELP condensates. The solution is sent to ICP-MS analysis. **(F)** Correlation between condensate fluorescence ratio (from C-SNARF-4) and ion concentration in the dense phase, as measured by ICP-MS. The results were fitted by the following formula: y = px + q, where p_salt, K_ = 1.417 M^−1^ (R^2^ = 0.8517) and p_salt, Na_= 1.739 M^−1^ (R^2^ = 0.8202). Data represent mean ± s.d. **(G)** Correlation between condensate fluorescence ratio and protein concentration in the dense phase, estimated from A_205_ absorbance using the calculated extinction coefficient of ELP and a fixed pathlength of 1 mm (NanoDrop 2000). The results were fitted by the following formula: y = px + q, p_protein, K_ = 3.742 M^−1^ (R^2^ = 0.9373) and p_protein, Na_ = 19.2 M^−1^ (R^2^ = 0.9297). Data represent mean ± s.d. **(H)** The pH capacity is defined as the ratio m_salt_ and m_protein_, representing the relative contribution of salt versus protein to pH modulation within condensates. Higher pH capacity indicates that pH changes are driven more by ionic partitioning than by protein concentration.

To induce the hydrophobicity effect, we added additional salt into the solution to salt out more water molecules from the protein backbone, which should create a more hydrophobic environment in the dense phase^28,44^. Interestingly, with increasing salt concentration, we observed a larger pH gradient between phases (**Figure 1C**). This pH gradient is mediated by a more basic interior pH and a stable dilute phase pH upon salt addition (**Figure S1**). This observation suggests that dense phase environment was altered by the additional salt, in line with hydrophobic effect driven by salt concentration-dependent water activity^45^.

To further alter the hydrophobic environment of the condensates, based on the Hofmeister effect of salts^46–48^, we implemented KCl, instead of NaCl as the salt to drive ELP phase separation. Compared to Na^+^, K^+^ is a more chaotropic ion in the Hofmeister series, thus possessing a stronger ability to salt out water molecules from the hydration shell of the protein backbone^48^. Indeed, we found that under the same ion concentration, the ELP condensates in KCl solution possessed a higher pH gradient compared to those in the NaCl solution (**Figure 1D**). Similarly, increasing salt concentration also increased the dense phase pH and enlarged the pH gradient between phases (**Figure S1**). The salt-dependent dense phase pH suggests that the salting out effect affects the interior ionic environment, which implies a role of hydrophobicity in affecting ion partitioning and water activity^49,50^.

### Decoupling the contribution of protein and salt on modulating condensate pH

As the salting-out effect should collectively affect ion and protein concentrations in the dense phase, we next sought out to decouple the contributions of dense phase ion concentration and protein concentration to condensate pH. To this end, based on different bulk ion concentration and identical total protein concentration, we generated condensates and employed the bulk phase separation assay^51^, and then separately analyzed the dense phase cation concentration using inductively-coupled mass spectrometry (ICP-MS) and the dense phase protein concentration (**Figure 1E**). We constructed two distinct correlation curves based on condensate pH against dense phase cation concentration and condensate pH against dense phase protein concentration using NaCl and KCl as the salts (**Figure 1F&G**). The slope of this correlation graph is defined as the pH capacity, which reflects the ability of ion or protein on modulating the condensate pH. By calculating the ratio between the pH capacity depending on ion and dense phase protein concentration based on NaCl and KCl, which allows us to compare the contribution of ion type onto the change of pH per unit change of protein concentration (**Figure 1H**), we identified that potassium ion contributed more significantly than sodium ion on regulating condensate pH. As the charge density of potassium ion is smaller than that of sodium ion^52^, the capability to generate a higher concentration of hydroxyl ions could not be explained through simple charge neutrality by oppositely charged ions. Thus, this effect might be caused by distinct ion effects on regulating water structure^53^, thus affecting the water activity. This finding suggests the importance of hydrophobicity of the interior environment on defining the effective autoionization of water molecules^54^, thus modulating condensate pH.

### Interior hydrophobicity-dependent pH gradient

To directly evaluate the role of condensate hydrophobicity in modulating its pH condition, we next adjusted the condensate microenvironment with organic small molecules, which bind with the available water in the dense phase^17^, thus altering the hydrophobicity of the interior of condensates. To this end, we added DMSO or 1,4-dioxane, which possesses distinct hydrogen bond capability. A previous work has shown that through analyzing the micropolarity of the ELP condensate^17^, compared with water, adding 1,4-dioxane created more polar interior environments, while adding DMSO created more non-polar interior environments (**Figure 2A**). Using the pH assay, we observed that compared to water, DMSO generated the most significant pH gradient while the 1,4-dioxane generated the least pH gradient between phases (**Figure 2B**). The capability of small molecule on regulating the pH gradient between phases follows the same trend as the effects of small molecules on regulating interior hydrophobicity.

**Figure 2.**
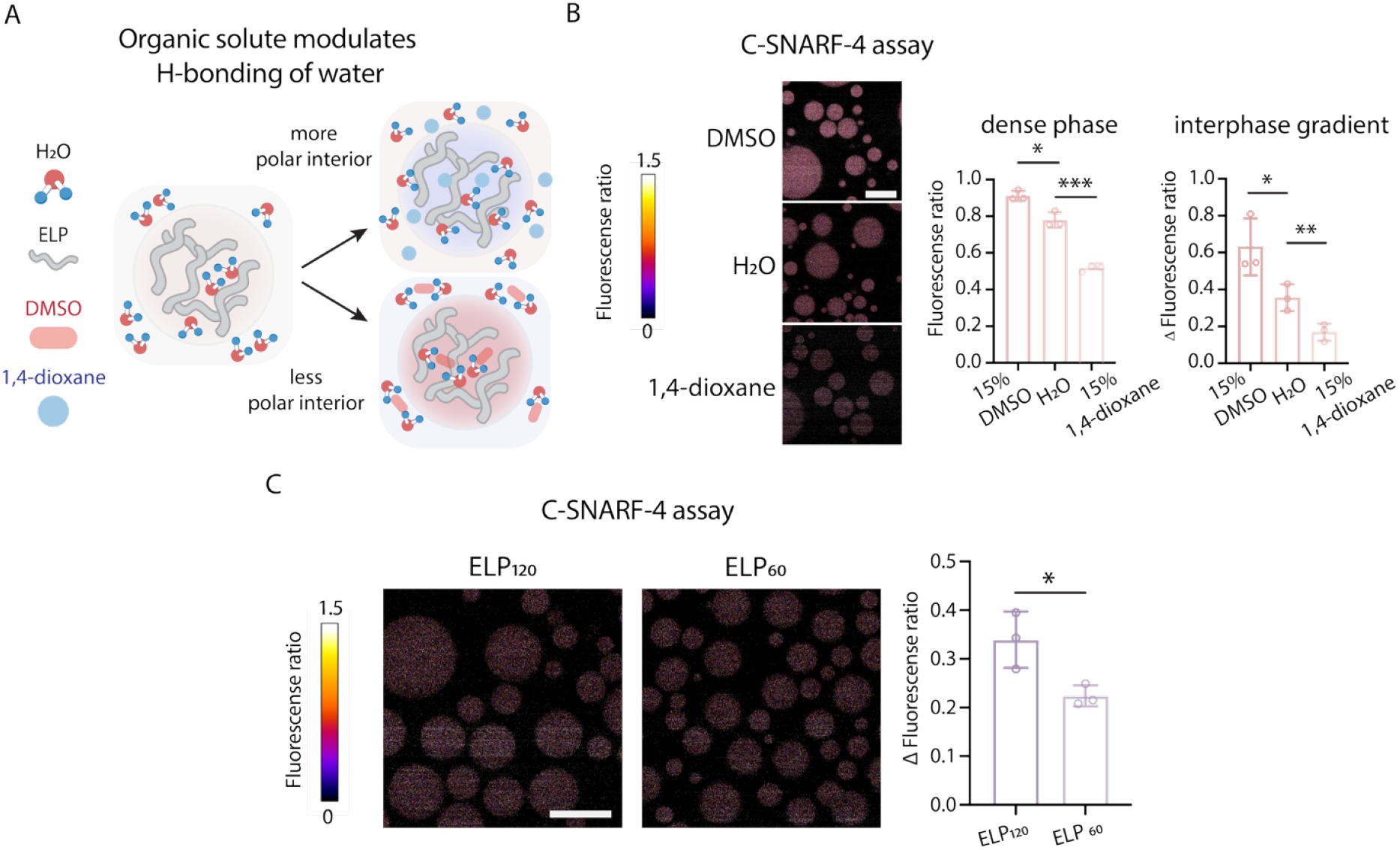
Interphase pH gradient is modulated by hydrophobicity effect. **(A)** Schematic of organic solute binds with water and modulate condensate polarity and hydrophobicity. **(B)** Representative C-SNARF-4 ratiometric images of ELP condensates formed in different solvent environments (15% DMSO, water, and 15% 1,4-dioxane), with constant salt type (KCl) and concentration (1 M). Pseudocolor scale represents pH, from acidic (blue) to basic (red). Δ Fluorescence ratio denotes the difference in fluorescence ratio between the dense and dilute phase. Data represent mean ± s.d. from n = 3 independent samples; >10 condensates of similar size were analyzed per sample. Statistical analysis was performed using a two-tailed t-test: *p < 0.05, **p < 0.01. Scale bar, 20 µm. **(C)** Representative C-SNARF-4 ratiometric images of ELP condensates formed from ELP variants of differing hydrophobicity. Pseudocolor scale represents pH, from acidic (blue) to basic (red). Δ Fluorescence ratio denotes the difference in fluorescence ratio between the dense and dilute phase. Data represent mean ± s.d. from n = 3 independent samples; >10 condensates of similar size were analyzed per sample. Statistical analysis was performed using a two-tailed t-test: *p < 0.05. Scale bar, 20 µm.

To further validate the role of hydrophobicity on encoding the dense phase pH, we reasoned that since the degree of polymerization directly scales with backbone hydrophobicity^32,55^, we next utilized ELP formed by 120 repeats of [Val-Pro-Gly-Val-Gly], twice the length of the ELP investigated in the previous sections. We found that indeed the dense phase pH of longer ELP is more basic than that of shorter ELP (**Figure 2C**). Collectively, these observations established a direct correlation between asymmetry in environmental hydrophobicity and pH gradient.

### Asymmetric intrinsic transfer free energy drives selective ion partitioning

The above data suggests that the basic interior environment within condensates could be explained by the cation partitioning effects by condensates. Based on the simplest theoretical explanation from electrochemical potential equilibria between phases, charge neutrality within one phase would require the partitioning of anions to neutralize the charge. However, the molecular mechanism by which cations partition into the dense phase is not clear.

To this end, we next performed all-atom molecular dynamics simulations to evaluate the molecular driving force underlying ion partitioning. We first constructed the dilute and the dense phases of ELP using all-atom molecular dynamics simulations, and calculated the intrinsic transfer free energies for some common cations (Li^+^, Na^+^, K^+^) and anions (F^-^, Cl^-^, Br^-^, I^-^) based on their intrinsic hydration free energies in the dense and the dilute phases ^56,57^: ΔΔ*G*_*I→II*_ = Δ*G*_*hyd,II*_ − Δ*G*_*hyd,I*_, where Δ*G*_*hyd,II*_, Δ*G*_*hyd,I*_ are the intrinsic hydration free energy of ions in the dense and the dilute phases, respectively. This simulation framework allows us to directly translate the effect of hydrophobicity in general into free energy of partitioning to understand the molecular driving force. Surprisingly, our simulations revealed that cations prefer the dense phase, exhibiting negative ΔΔ*G*_*I→II*_ values, whereas anions favor the dilute phase with positive ΔΔ*G*_*I→II*_ values (**Figure 3A**). By analyzing the interaction strength between the carbonyl oxygen of the amide bond on the protein backbone and different cations, as well as interactions between the amide nitrogen and various anions, we found that this asymmetry in intrinsic transfer free energy is likely attributed to distinct interaction propensity between ions and protein backbone (**Figure 3B**). Further free energy analysis demonstrates that the intrinsic transfer free energy is mainly contributed by enthalpy (**Table S1**). This finding indicates that a higher protein concentration within the condensate will lead to more favorable transfer free energy for cations (**Figure S2**), which is consistent with the positive correlation between condensate protein concentration and ion partitioning observed in experiments.

**Figure 3.**
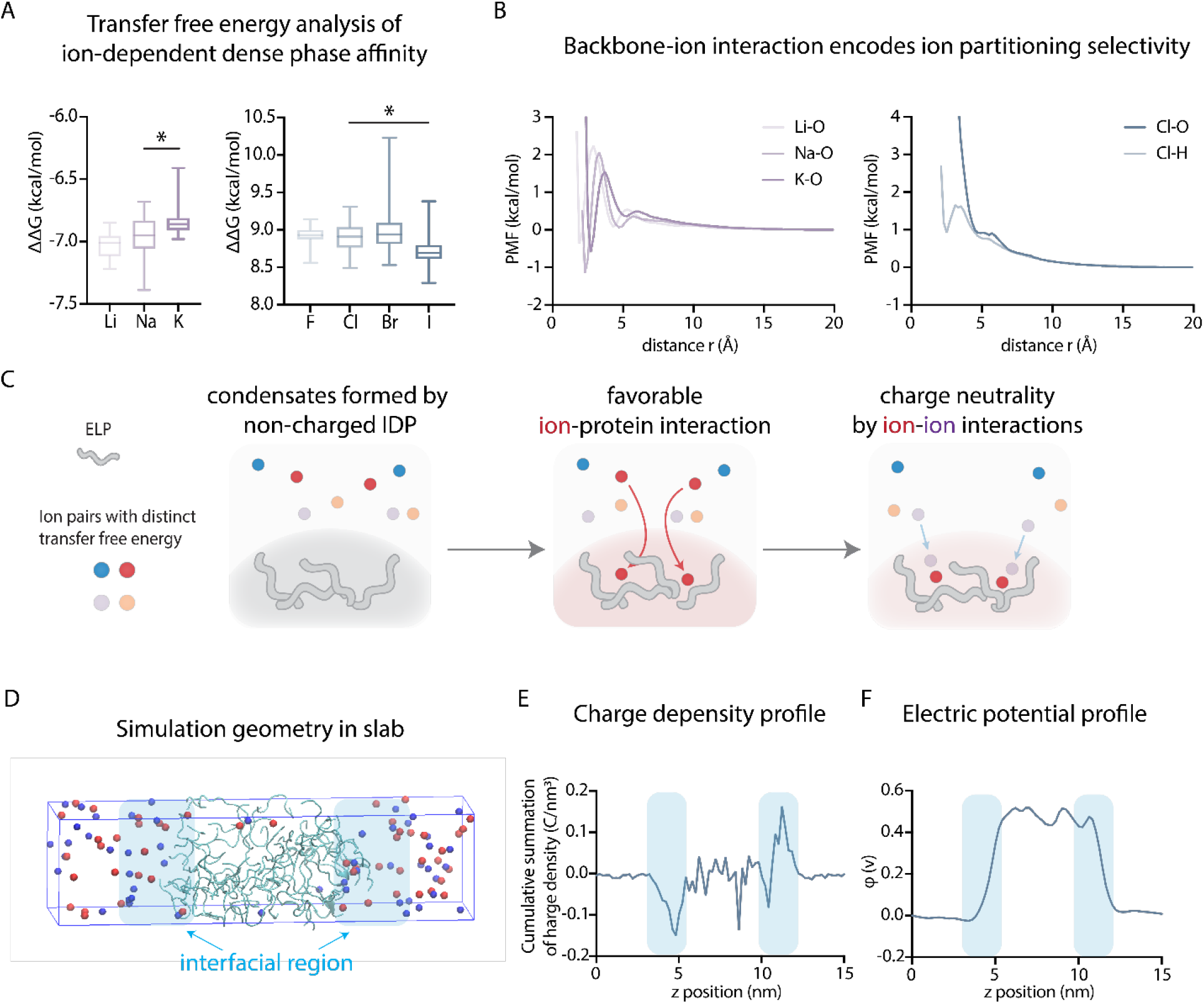
Molecular dynamic simulations reveal selective ion partitioning model and charge and potential distributions of ELP condensates. **(A)** Asymmetric intrinsic transfer free energy ΔΔ*G* for cations and anions. For each ion, 15 independent free energy calculations are performed to calculate ΔΔ*G*. **(B)** The potential of mean forces (PMFs) between cations (Li^+^, Na^+^, K+) and the backbone carboxyl oxygen atoms (left panel), between anion Cl^-^ and the backbone carboxyl oxygen atoms (right panel), and between anion Cl^-^ and the backbone amide hydrogen atoms (right panel). **(C)** Schematic of the proposed ion partitioning model. The “active” partitioning of cations due to the thermodynamically favorable intrinsic transfer free energy drives the “passive” partitioning of anions. Cations and anions are represented by red/orange and blue/purple spheres, respectively. **(D)** A snapshot of the all-atom molecular dynamics simulation visualized by the VMD software. A slab of dense phase coexists with two regions of the dilute phases. Cations and anions are represented by blue and red spheres, respectively. Peptides are shown by the ‘NewCartoon’ representation. Water molecules are not shown for clarity. The interfacial regions are highlighted by light blue. **(E)** The cumulative summation of net charge density profile along the z axis, *C*(*ρ*(*z′*), *z*) calculated as the integral of the net charge density along the z-axis of the simulation box (*ρ*(*z′*)) from all-atom molecular dynamics simulations.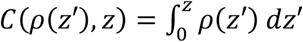. The trough on the left interface and the peak on the right interface indicates the existence of electric double layers at these two interfaces. **(F)** The electrostatic potential profile *Φ* along the z-axis of the simulation box calculated from all-atom molecular dynamics simulations.

Mechanistically, these results suggest that during the initial phase separation process, cations first partition into the dense phase due to favorable intrinsic transfer free energies. Subsequently, anions are passively attracted to the dense phase to neutralize the net charge. Thus, we propose that the cation partitioning resembles an “active process”, while anion partitioning can be considered a “passive process” (**Figure 3C**). This mechanism can be placed within the broader context of phase separation. The efficiency of the “active” cation partitioning is directly dependent on the number of available protein backbone carbonyls— in other words, on the protein concentration within the condensate. It is well-established that this protein concentration itself is governed by the overall physicochemical environment, including the salting-out effect and the hydrophobicity of the condensate’s interior. Thus, these results indicate that the asymmetry in the hydrophobicity leads to distinct intrinsic transfer free energies for ions, resulting in distinct partitioning functions of ions into the dense phase, which ultimately sets up the pH gradient.

### The formation of interfacial potential and electric double layer

Previous studies suggested that an established potential gradient between phases manifests as the formation of an electric double layer at the liquid-liquid interface of condensates^6,7^. This interface is defined as the Gibbs dividing plane^22^. To investigate this, we analyzed the coexisting dilute and dense phases of ELPs under all-atom molecular dynamics simulations with a slab geometry^58^. In these simulations, the dense phase occupies the central region of the simulation box flanked by the dilute phase on both sides (**Figure 3D**).

By analyzing the cumulative net charges along the slab’s long axis, we identified a region with net negative charges in the dilute-phase side of the left interface (**Figure 3E**), and the cumulative net charges transition to 0 when entering the dense phase, suggesting a region with net positive charges in the dense-phase side of the interface. A similar feature was observed near the right interface. This charge distribution suggests the presence of an electric double layer at the interface of ELP condensates.

Based on the ion distribution profile, we next calculated the electric potential distribution ^59^and found a higher electric potential in the dense phase compared to the dilute phase (**Figure 3F**). This elevated electric potential counteracts the favorable intrinsic transfer free energy for cations, explaining their low equilibrium concentration in the dense phase. The higher electric potential in the dense phase also supports the higher dense phase pH as the Galvani potential gradient is positively correlated with the pH gradient between the dense and the dilute phase as identified previously^8^, 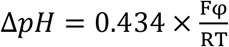, where *φ* represents the theoretical Galvani potential between phases. Then, by analyzing the localized interfacial potential gradient, we calculated the strength of interfacial electric field, which is at a level of 10^8^ V/m. This averaged interfacial electric field is strong enough to potentially drive electrochemical reactions as observed previously at diverse liquid-liquid, liquid-air interfaces^7,60–62^.

### Selective ion effect encodes the directionality of the pH gradient

We next explored whether anions could serve as a source to modulate the dense phase pH. To this end, we compared the dense phase pH of condensates formed by KCl, KI and KBr, which also possess distinct free energy of transfer as calculated through our molecular dynamics simulation (**Figure S3**). Experimentally, we found that indeed, ELP condensates formed in KI and KBr possessed an apparently lower pH than the ELP condensates formed in KCl (**Figure 4A**). Thus, we compared the ΔΔG of Cl^-^, Br^-^, and I^-^, and found that I^-^ has a less positive value (**Figure S3**), suggesting a stronger ability to partition into the condensates. Due to the competition between I^-^and OH^-^, less OH^-^ would partition into the dense phase to mediate phase charge neutrality, resulting in a lower pH. These observations suggest that the competition between ions in the solution system can directly alter the electrochemical properties of condensates.

**Figure 4.**
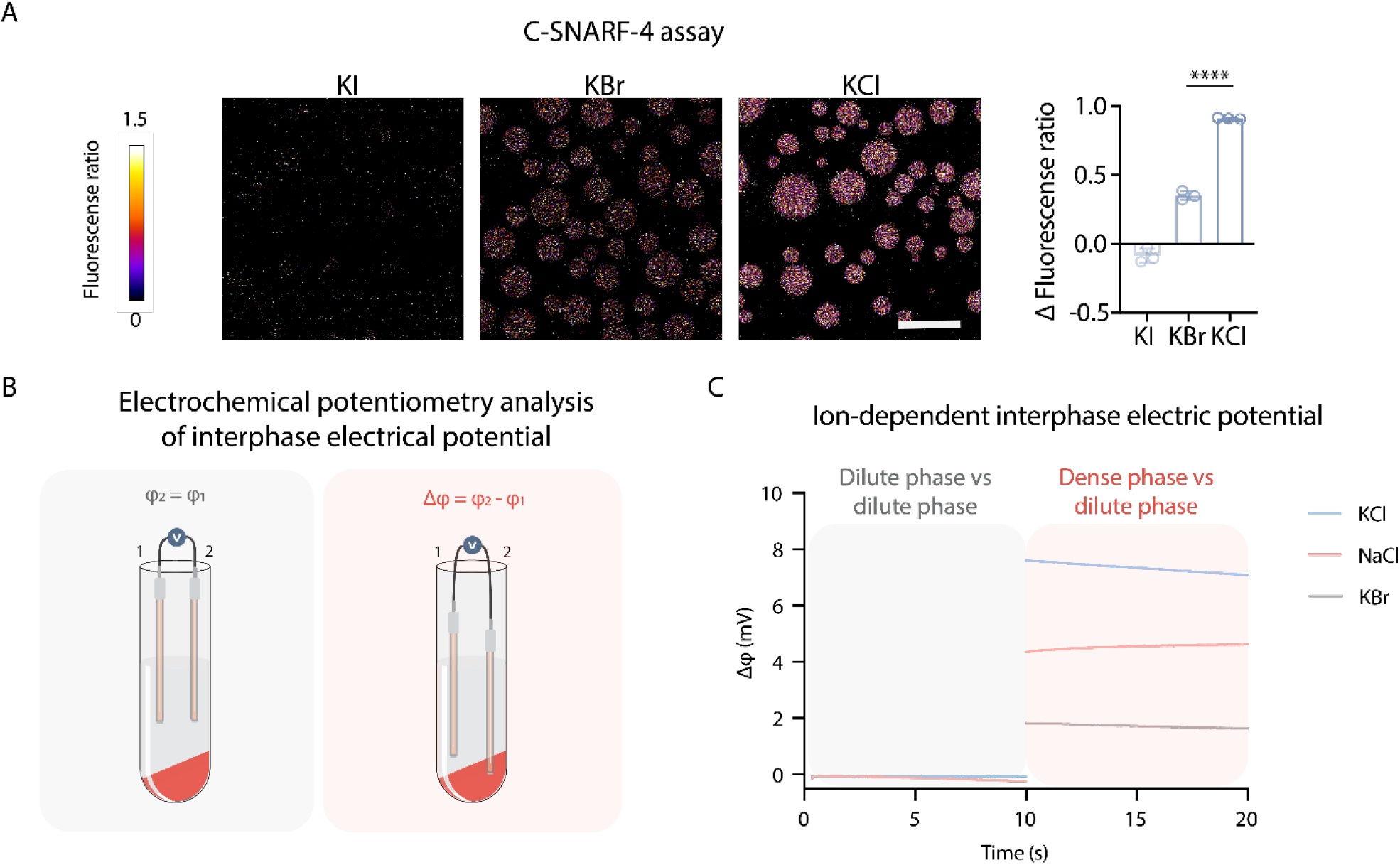
Direct analysis of hydrophobicity-dependent interphase electric potential. **(A)** Representative C-SNARF-4 ratiometric fluorescence images of ELP condensates formed in 1.5 M solutions of KI, KBr, or KCl. Pseudocolor scale represents pH, from acidic (blue) to basic (red). Δ Fluorescence ratio denotes the difference in fluorescence ratio between the dense and dilute phases. Data represent mean ± s.d. from n = 3 independent samples; >10 condensates of similar size were analyzed per sample. Statistical analysis was performed using a two-tailed t-test: ****p < 0.0001. Scale bar, 20 µm. **(B)** Schematic of electrochemical potentiometry analysis of interphase electrical potential difference between the dense and dilute phase. High-speed centrifugation separates the two phases, and the two Ag/AgCl electrodes measure interphase electrical potential difference. **(C)** Measurements of the interphase electric potential difference for a condensate sample prepared in different salt. Open-circuit potential measurements (10 s) were performed with the two-electrode set-up. The colored backgrounds correspond to distinct set-up showed in

### Direct analysis of interphase electric potentials

With these distinct ion-dependent charging behaviors of ELP condensates, we next implemented our customized electrochemical potentiometry to directly analyze the interphase electric potentials (**Figure 4B**). This setup measures the open circuit potential between two identical Ag/AgCl electrodes, which are respectively inserted into the dilute and the dense phases, so that the differences in liquid-junction potential between the solution phase and the electrode would be reflected as the interphase potential^8^. We found that consistent with the pH gradients across ELP condensates induced by NaCl, KCl, and KBr, the interphase potential of these condensates exhibited the same trend (**Figure 4C**). This experiment demonstrates the distinct capability of salts on charging the ELP condensates.

### Electric potential gradient regulates the partitioning/exclusion of charged small molecules

An electric potential gradient acts as a driving force modulating the transport of charged small molecules based on Nernst equilibrium. Thus, we reasoned that when condensates are potentiated differently by different types of ions, then the directionality of the electric potential gradient might define the partitioning of small molecules that are charged differently (**Figure 5A**). This might serve as the underlying factor encoding condensate reaction selectivity. To this end, we employed condensates charged by NaCl, KCl and KBr, which showed distinct electric potential gradient, and evaluated the partitioning of charged fluorescent molecules, including cationic Rhodamine 6G and anionic Alexa 647 NHS ester (**Figure 5B**). As expected, we observed that the partitioning ratio of Rhodamine 6G or Alexa 647 follows the same trend of the electric potential gradient of ELP condensates charged by different salts. This observation suggests that the electric potential of condensates could encode selectivity of small molecules partitioning based on their charge conditions, thereby shaping their reactivities.

**Figure 5.**
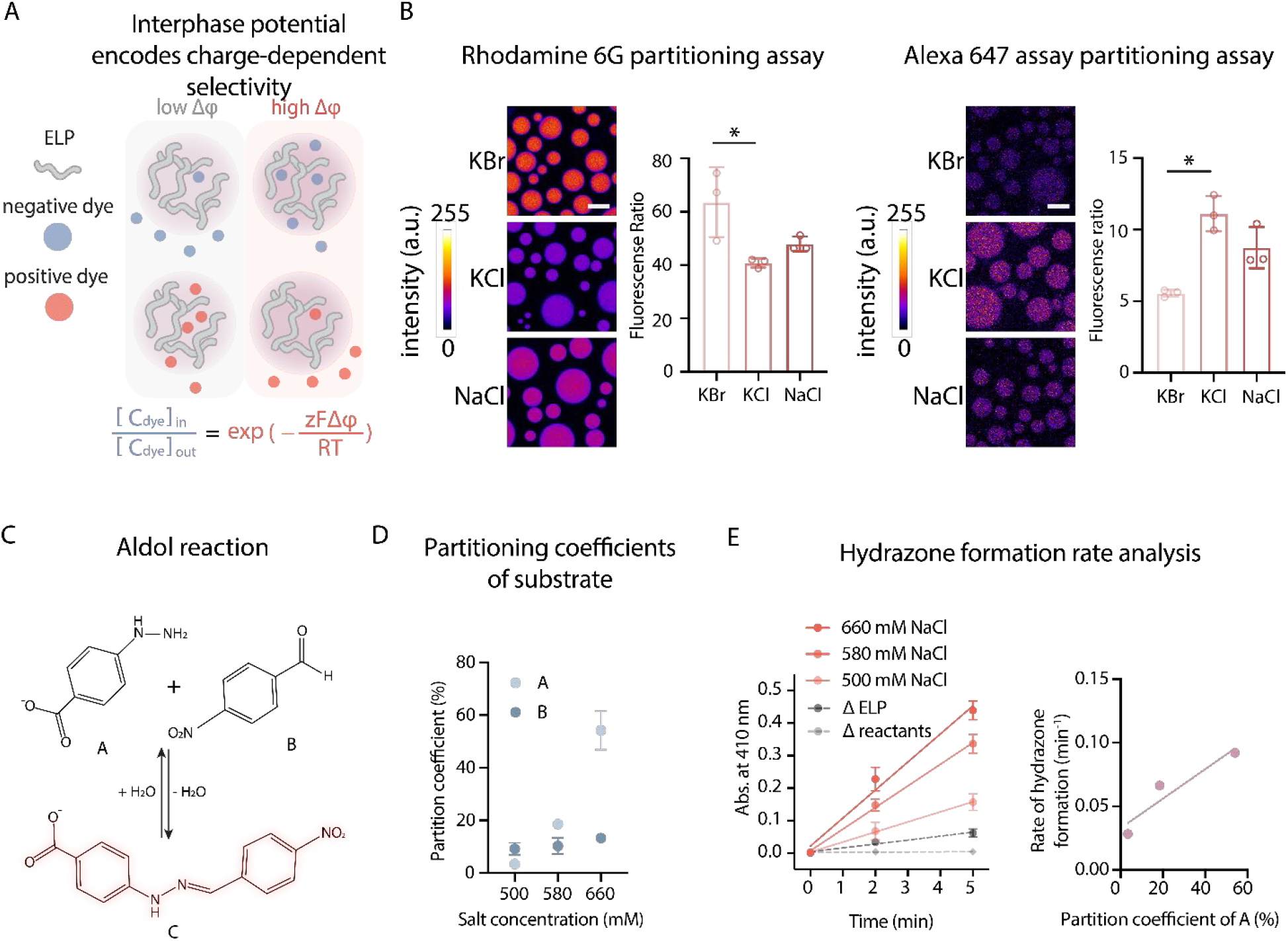
Condensate electric potential encodes selectivity and modulates reaction kinetics for non-enzymatic chemistry of condensates. **(A)** Schematic illustration of how interfacial electric fields regulate condensate ability to partition charged species. **(B)** Evaluation of negative charge partitioning using Alexa Fluor 647 NHS ester dye and positive charge partitioning using Rhodamind 6G dye in ELP condensates formed in 1 M solutions of KCl, KBr, or NaCl. Fluorescence ratio denotes the intensity in the dense phase divided by that in the dilute phase. Data represent mean ± s.d. from *n* = 3 independent samples; >10 condensates of similar size were analyzed per condition. *p < 0.05. Scale bar, 10 µm. **(C)** Reaction scheme of hydrazone formation between 4-nitrobenzaldehyde and 4-hydrazinobenzoic acid. The product absorbs at λ_max_ ≈ 410 nm. **(D)** Partition coefficient of reactant A and B into ELP condensates formed at 500, 580, and 660 mM NaCl. Data represent mean ± s.d. from *n* = 3 independent samples. **(E)** Hydrazone formation reaction for ELP condensates formed at 500, 580, and 660 mM NaCl compared to controls that have either no reactants or no ELP condensates. Initial rate of hydrazone formation is plotted against partition coefficient of reactant A, and the results were fitted by the following formula: y = px + b, where p = 0.001159 ± 0.000162, which is significantly different from 0. Data represent mean ± s.d. from *n* = 3 independent samples.

### Electrochemically active condensates drive potential-dependent non-enzymatic reactions

We next evaluated whether the electrochemically active ELP condensates could power electrochemical reactions. Using the previously demonstrated resazurin and PY1 assay^6,63^, we found that the charging the ELP condensates through hydrophobic effects can activate their electrochemical functions (**Figure S4**). To understand whether the electrochemical properties of condensates could govern the reaction selectivity, we implemented a dehydration reaction as demonstrated previously^15,17^. The previous work showed that the ELP condensates could power non-enzymatic dehydration reaction to form hydrazone^17^. Using this reaction as the model reaction, we next examined the ability of ELP condensates with distinct interphase potentials to promote this reaction (**Figure 5C**). We chose to vary the concentration of a single salt rather than comparing different salts when probing hydrazone formation within condensates. Changing salt identity would introduce additional confounding variables, particularly in the context of a chemical reaction. Distinct ions can differentially alter the condensate microenvironment, such as specific ion–protein interactions, therefore making it difficult to disentangle effects due to partitioning from those due to ion chemistry. By instead adjusting the concentration of the same salt, we were able to tune condensate partitioning capacity while excluding ion-type dependent chemistry, thereby ensuring that differences in reaction outcome primarily reflect the changes in reactant partitioning rather than ion-specific effects.

We first verified that the interphase potential-dependent partitioning effect of charged small molecules could be regulated by salt concentration (**Figure S5**). We next utilized the intrinsic fluorescence of reactant A (4-hydrazionobenzoic acid) and reactant B (4-nitrobenzaldehyde) to evaluate their partitioning functions in the condensates charged by distinct salt concentrations (**Figure 5D**). To confirm the absorbance peak of product C, we performed *λ*-scan and found peak at 410 nm (**Figure S6A**). We observed that the partitioning of the negatively charged reactant A depends on the interphase potential of condensates, while the non-charged reactant B showed limited differences between differently charged condensates. The *λ*_max_ for reactant A is at 230 nm, which is close to 205 nm, the wavelength we used to measure protein concentration. This overlap raises the possibility that our measurements could be influenced by changes in protein concentration across conditions. However, protein concentration only increased about 8-fold, whereas reactant A absorbance increased about 18-fold. This disparity indicates that the observed signal cannot be explained by protein alone and demonstrates that reactant A indeed partitions more strongly into the dense phase as bulk salt concentration increases. By introducing both reactant A and B into different condensate systems and monitoring the formation of the product via its native absorbance, we observed that condensates with the highest interphase potential showed the fastest reaction kinetics (**Figure 5E**). Using confocal microscopy imaging to analyze the condensate solution, we confirmed that the size and the number of condensates were similar between samples (**Figure S6B&C**). By plotting reaction rate against partition coefficient of reactant A, we found positive linear correlation with a R-square value of 0.8802 (**Figure 5F**). These observations suggest that the asymmetry in hydrophobicity-dependent charging effects encodes the capability of condensates to power non-enzymatic reactions and also govern reaction selectivity.

## Discussion

The origin of hydrophobicity lies in the electromagnetic organization of water and related solvents. In this view, solvent polarity is not merely a background property but an active determinant of electrochemical environments. When two phases differ in hydrophobicity, their asymmetric solvent structuring imprints an interphase electrical potential gradient. Our findings place this principle within biomolecular condensates, showing that ion partitioning magnifies solvent asymmetry to yield electric potential gradients between phases. This perspective reframes condensates as electrochemically active entities: their microenvironments are shaped not only by molecular composition but by solvent polarity gradients^17,27,64^. These findings suggest that water environments serve as the most fundamental feature of condensates governing their electrochemical properties.

With all-atom molecular dynamics simulations, which provide an essential bridge from macroscopic experimental observations to the underlying molecular mechanism, we uncovered the driving forces for the interphase potential at the atomistic level. We directly calculated the intrinsic transfer free energy (ΔΔG) for individual ions moving into the dense phase, which provided two critical insights. First, it revealed an apparent energetic preference for cations to enter the condensate while anions were repelled. This elucidated the origin of this asymmetry, namely a strong, enthalpically favorable interaction between the cations and the carbonyl oxygen atoms of the protein backbone. This phenomenon should be generalizable to any protein condensate. Second, the simulations directly revealed the electric-double-layer-like structure at the condensate interfacial region, which leads to the interfacial potential. Part of this potential is simply set up by the water dipole orientation, which also suggests its generality. Taken together, the simulations provided the high-resolution, mechanistic evidence necessary to propose the “active/passive” partitioning model and provided direct evidence to the molecular origin of the interphase potential at the atomistic level.

How is the electrochemical equilibrium being realized in such a complex biomolecular two-phase system? There is a complex interplay between interphase potential, ion partitioning, internal environment and interfacial molecular organization: 1) the interphase potential accompanied with an electric field introduces one more term in the chemical potential of ions, which affects the phase equilibrium of ions and ion partitioning to condensates^21^; 2) ion partitioning modulates the internal environment of condensates, such as modulating the interior environmental polarity^27,64^; 3) the internal environment of condensates defines the transfer free energy of ions, thereby in turn modulating ion partitioning; 4) the formation of double layer structure at the liquid-liquid interface affects the ion distribution in the dense phase and the dilute phase and the molecular organization at the surface of condensates further modulate local ion equilibrium^11^. As such, interplay between different factors and their relative thermodynamic and kinetic driving force should collectively define the pseudo-equilibrium of electrochemical potential of condensates, which encode condensate electrochemistry.

Through experiments, we revealed how condensates could simply exploit the principle of Nernst equilibrium to modulate small molecule distribution, which in turn determines the functional selectivity of condensates. This aspect suggests that defining the electrochemical potential profiles of different condensates will provide the foundation to understand the foundation for how condensate functions could be regulated by metabolite distributions, another important frontier in condensate science^65^. Our study provides a water-centered view of electrochemistry of condensates, which suggests the generality of electric potential gradients and electrochemical functions in condensate systems.

## Acknowledgements

We appreciate the discussion with Siqin Cao and Jing Huang from University of Wisconsin-Madison and Westlake University. Y.D. acknowledge the funding support from the McKelvey School of Engineering and the Center for Biomolecular Condensates.

## Author Contributions

Y.D. generated the idea and devised the study. L.Y., X.Z. and Y.D. designed the experiments. L.Y., X.Z., W.Y. performed the experiments. Y.D. and X.Z. wrote the initial draft of the manuscript and all the authors contributed to the revision of the manuscript. Y.D. acquired funding.

## Declaration of Interests

The authors declare no competing interests.

## Inclusion and Diversity

We support inclusive, diverse and equitable conduct of research.

## Notes

### Competing Interest Statement

The authors have declared no competing interest.

