## Supplementary material for "Asymmetry in hydrophobicity induces electric potential in non-charged protein condensates": 10-15-2025 Supplementary Information.pdf

### Theoretical analysis procedures

Calculation of intrinsic transfer free energies ($\Delta\Delta G$) from all-atom molecular dynamics simulations

Both the intrinsic transfer free energy ($\Delta\Delta G$) and interphase potential were calculated using all-atom molecular dynamics simulations using the Gromacs 2024.2 package [1] on the MGPU cluster at the High Performance Computing Center (HPCCC) of Hong Kong Baptist University. $\Delta\Delta G$ for ions was determined by comparing the intrinsic hydration free energy of ions in the bulk water ($\Delta G_{hyd,I}$) to that in a dense-phase mimic system containing peptides ($\Delta G_{hyd,II}$). The hydration free energies were calculated using the Bennett Acceptance Ratio (BAR) method [2] implemented in Gromacs. The ions are grown using two different coupling parameters $\lambda_{vdw}$ and $\lambda_{ele}$. We used the following combinations for these two coupling parameters: $[\lambda_{vdw}, \lambda_{ele}] =$ [0, 0], [0.1, 0], [0.2, 0], [0.3, 0], [0.4, 0], [0.5, 0], [0.6, 0], [0.7, 0], [0.8, 0], [0.9, 0], [1.0, 0], [1.0, 0.1], [1.0, 0.2], [1.0, 0.3], [1.0, 0.4], [1.0, 0.5], [1.0, 0.6], [1.0, 0.7], [1.0, 0.8], [1.0, 0.9], [1.0,1.0].

A cubic simulation box with a length of ~ 4 nm was used for both $\Delta G_{hyd,I}$ and $\Delta G_{hyd,II}$. For $\Delta G_{hyd,I}$, 10 ns long simulations were performed for each $\lambda$ pair, with the last 9 ns for free energy calculations. For each ion, 10 independent free energy calculations were performed. For the calculation of the $\Delta G_{hyd,II}$, 20 ns long simulations were performed for each $\lambda$ pair, with the last 19 ns used for free energy calculations, and 15 independent free energy calculations were performed for each ion.

The dense phase was modeled as a cubic water box with a length of ~4nm containing 30 chains of ACE-$(VPGVG)_2$-NME sequences, achieving a solution density of ~1.18 g/cm$^3$. Initial peptide conformations were generated from the excluded volume simulations using the CAMPARI molecular simulation software (https://campari.sourceforge.net/). Then, the peptides were solvated in the water box, followed by an energy minimization. Next the system was equilibrated using the NVT ensemble for 500 ps and NPT ensemble for 400 ps. A 1 µs production simulation was then performed, with the final frame used for the free energy calculation.

Simulation details and forcefield considerations

The Amber ff14sb forcefields parameters [3] were used for the peptides and tip3p water model [4] was used for the simulations. Parameters developed by Joung and Cheatham [5] were used for ions. All the simulations were performed under the NPT ensemble. The temperature was kept at 298 K using the V-rescale thermostat [6] with a coupling time constant of 2 ps. The pressure was maintained at 1 bar a coupling time constant of 2.0 ps using the c-rescale method [7]. Cutoffs for short-range electrostatic potential and van der Waals potential were set to 1.2 nm and 1.1 nm, respectively. Periodic boundary conditions were applied and long-rang electrostatic interactions were calculated using the Particle Mesh Ewald (PME) algorithm [8,9]. The LINCS algorithm [10] was used to constrain all bonds with H-atoms. A leap-frog stochastic dynamics integrator, sd option for integrator in Gromacs, with an integration step 2 fs was used.

To investigate the influence of forcefields, we also calculated $\Delta G_{hyd,II}$ and $\Delta G_{hyd,I}$ using the amber03ws forcefields [11] with tip4p/2005 water model [12]. While small differences were observed in the absolute values, the relative trends among ions and the asymmetry between cations and anions remained consistent (**Fig. S3**).

<u>Entropy and enthalpy decomposition of $\Delta\Delta G$</u>

To decompose $\Delta\Delta G$ into enthalpy and entropy components, the simulation time for each pair of $\lambda$ values was extended to 100 ns with potential energy data saved every 200 ps. 10 and 5 independent free energy calculations were performed for $\Delta G_{hyd,II}$ and $\Delta G_{hyd,I}$, respectively. Therefore, a total of 31.5 μs simulations were performed for each ion. A lower peptide concentration, modeled with 20 chains of ACE-$(VPGVG)_2$-NME sequences in a $4 \times 4 \times 4\ nm^3$ water box, was used for energy decomposition. The method proposed by Wyczalkowski et al [13] was used for estimating the entropy and enthalpy contributions.

<u>Calculation of potential of mean forces (PMF) between ions and peptide backbone</u>

5 independent simulations containing one ACE-$(VPGVG)_2$-NME peptide and 250 mM ions in a simulation box with a length of ~ 4 nm were performed. Each simulation is 1 μs long. Then the distance distribution between ion and peptide backbone atom $P(r)$ was calculated, and the PMF was calculated as $W(r) = -\ln\frac{P(r)}{4\pi r^2} + C$, where $C$ is a constant shifting the value of $W(r = 2nm)$ to 0.

<u>Calculation of interphase potential from all-atom simulations with a slab geometry</u>

To construct a simulation box with coexisting dense and dilute phases, we first performed simulations using a box size of ~ $4 \times 4 \times 8\ nm^3$ with 60 chains of ACE-$(VPGVG)_2$-NME peptides to model the dense phase. The simulation protocol of this dense phase was identical to that described earlier. Initial peptide conformations were obtained using CAMAPRI and a 1 μs production simulation was performed to equilibrate the system. Then, the z direction of the last frame was extended to 16 nm, and water molecules were added to fill the simulation box. Next, 320 mM NaCl were added to the system. The system was equilibrated using the NVT ensemble for 500 ps, and NPT ensemble for 200 ps. 3 μs production simulation were performed, with the last 2.8 μs to calculate the interphase potential. The amber03ws forcefields [11] with tip4p/2005 water model [12] were used for these simulations. All the other simulation details are identical to that described earlier. Then the interphase potential was calculated using the 'gmx potential' program implemented in Gromacs, which derives the electric potential by integrating the net charge density profile [14].

**Experimental procedures**

Plasmids, strains, kit, and media

All DNA fragments encoding the proteins of interest were synthesized by Integrated DNA Technologies (IDT). Plasmid maps were generated using SnapGene software. *E. coli* strains BL21(DE3) and DH5α, used for protein expression and cloning, respectively, were purchased from New England Biolabs.

2xYT medium supplemented with 0.4% (w/v) glucose and corresponding antibiotics was used for routine culturing. For in vivo protein expression experiments, M9 minimal medium was supplemented with 2 mM $MgSO_4$, 100 µM $CaCl_2$, 0.4% (w/v) glucose, and 0.2% (w/v) casamino acids. Unless otherwise specified, all cultures were grown at 37 °C with constant shaking at 250 rpm in an orbital shaker.

Protein expression and purification

**Expression:** ELP constructs were expressed in *E. coli* BL21 (DE3) cells transformed with the expression plasmid. Single colonies were cultured overnight at 37 °C in 2xYT medium supplemented with 45 mg/L kanamycin. This starter culture was diluted 1:100 into 1 L of fresh 2xYT-kanamycin media and incubated at 37 °C until reaching an $OD_{600}$ of ~0.5. Protein expression was induced by adding 0.5 mM IPTG, followed by 24 h incubation at 30 °C with shaking at 250 rpm.

**Purification of ELP (sequence: $[VPGVG]_{60}$-VPGY, $[VPGVG]_{120}$-VPGY):** Purification leveraged ELP's temperature-responsive solubility (LCST behavior). Harvested cell pellets from 1 L cultures were collected by centrifuging at 4,000 × g for 10 min at 4 °C and resuspended in 30 mL of ice-cold PBS (1×). Cells were lysed by sonication (2 s on/2 s off, 30% amplitude, 4 min total) in an ice bath. To degrade nucleic acids, 500 U of Benzonase (MilliporeSigma) was added, and the mixture was incubated at 4 °C for 1 h. The lysate was cleared by centrifugation at 20,000 × g for 30 min at 4 °C, and the supernatant was adjusted to 1 M NaCl and incubated at 55 °C for 3 h to trigger ELP coacervation. Precipitated protein was pelleted by centrifugation at 20,000 × g for 30 min at 40 °C, redissolved in cold PBS (1×), and incubated at 4 °C for at least 6 h. This hot/cold phase transition cycle was repeated three times for further purification. Protein purity was assessed via SDS-PAGE using Any kD Mini-PROTEAN TGX gels (Bio-Rad), revealing >95% purity after Coomassie staining. Final samples were dialyzed extensively against Milli-Q water at 4 °C with at least three buffer exchanges. Samples were frozen at -80 °C and lyophilized for 72 h (Labconco FreeZone 2.5) prior to storage at -20 °C.

Sample preparation for confocal microscopy characterization

Lyophilized ELP protein was weighed and dissolved in Milli-Q water to prepare protein-containing solutions for imaging experiments. Salts were introduced by diluting from concentrated stocks into protein-containing solution to achieve desired final concentrations. For all comparative assays, the ELP concentration was maintained at 1 mM.

To minimize confounding size effects, only condensates of comparable diameters were selected for quantitative image analysis.

C-SNARF-4 assay to evaluate relative pH values

To assess the pH environment within condensates, the ratiometric dye C-SNARF-4 (Thermo Fisher Scientific) was added from a 200 μM stock solution to a final concentration of 2 μM. Samples were incubated at room temperature in PhenoPlate™ 384-well microplates (Revvity) for 10 min to allow dye equilibration. Confocal fluorescence images were acquired using a Leica Stellaris 8 Falcon equipped with an oil-immersion objective. The dye was excited at 490 nm, and emission was collected in two channels: 540-590 nm (channel 1) and 610-660 nm (channel 2). Regions of interest (ROIs) encompassing entire condensates were manually selected using ImageJ. Mean pixel intensities in each channel were extracted, and the ratio of channel 2 to channel 1 was used as a relative indicator of local pH.

To determine the signal corresponding to the dilute phase, the same condensate-containing samples were centrifuged at room temperature at 20,000 × g to separate the dense and dilute phases. The dye concentration of the recovered dilute phase was adjusted to a final concentration of 20 μM to ensure robust detection, as signal in dye-free dilute phase was often below the detection threshold. Because the dye is ratiometric, this higher concentration is appropriate for relative comparison. Quantification of condensate pH shift was performed by subtracting the mean ratio of the dilute phase from that of the dense phase.

ICP-MS analysis of condensate solution

To assess ionic partitioning between the condensate and dilute phases, samples were centrifuged at room temperature at 20,000 × g to separate the two phases. Immediately after centrifugation, the supernatant was carefully removed, and 1 μL of the dense phase was transferred using a positive-displacement pipette into 100 μL of Milli-Q water to minimize errors during extraction of dense phase pellet. Both the dilute and dense phase fractions were acidified with trace-metal grade nitric acid to a final concentration of 1% (v/v) prior to analysis. Inductively coupled plasma mass spectrometry (ICP-MS) was performed using a NexION 2000 instrument (PerkinElmer) operating in collision cell mode under helium gas. Calibration was performed using a mixed-element standard containing sodium, potassium, calcium, and magnesium. All sample processing was carried out using metal-free tubes and ultrapure reagents to minimize contamination. Final ion concentrations were back-calculated from ICP-MS readouts using the known dilution factors for each phase.

Protein concentration analysis for the dense phase of condensate solution

To determine the protein concentration within the dense phase of ELP condensates, dense phase and dilute phase were separated as described in the previous sections. Absorbance at 205 nm ($A_{205}$) was measured using a NanoDrop 2000 spectrophotometer (Thermo Fisher Scientific). The extinction coefficient was estimated using a sequence-based method previously described[15], which accounts for peptide backbone absorbance in the absence of aromatic residues. Extinction coefficient at 205 nm for $[VPGVG]_{60}$ is 831220 $M^{-1}$ $cm^{-1}$. All measurements were performed using a fixed pathlength of 1 mm, and protein concentration was calculated using the Beer-Lambert Law. Measurements were only considered in the linear range of the spectrophotometer ($A_{205}$ between 0.5-1.5), and blank (water) was subtracted to account for background absorbance.

Resazurin assay for assessing reducing capacity

To evaluate the reducing capability of condensates, resazurin (Thermo Fisher Scientific) was added to a final concentration of 1 μM. Samples were incubated for 10 min at room temperature in PhenoPlate™ 384-well microplates prior to imaging. Confocal fluorescence images were captured using a Leica Stellaris 8 Falcon with an oil-immersion objective, using excitation at 532 nm and collecting emission from 540-590 nm. Images were acquired at two time points: immediately after incubation (0 h) and after 1 h. ROIs were manually selected around individual condensates using ImageJ, and mean pixel intensities were quantified.

PY1 assay for the characterization of $H_2O_2$ production

Peroxy Yellow 1 (PY1; Thermo Fisher Scientific) was used to quantify $H_2O_2$ generation within condensates. A 1 mM dimethyl sulfoxide (DMSO) stock solution was diluted into samples to achieve a final concentration of 20 μM. Samples were incubated for 10 min at room temperature in PhenoPlate™ 384-well microplates prior to imaging. Fluorescence imaging was performed on a Leica Stellaris 8 Falcon with an oil-immersion objective. The dye was excited at 514 nm, and emission was collected from 538-558 nm. Images were acquired at two time points: immediately after incubation (0 h) and after 1 h. ROIs were manually selected around individual condensates using ImageJ, and mean pixel intensities were quantified.

Electrochemical potentiometry for interphase electric potential difference

Phase electric potential difference measurements were carried out in a 40-mL high-speed polycarbonate centrifuge tube (Thermo Scientific). 15 mL of solution containing ELP condensates was centrifuged at 40 °C at 20,000 × g for 30 min to separate the phases. Protein concentration was 0.5 mM and salt concentration was 1 M. Two identical Ag/AgCl reference electrodes were used to measure the potential difference between the two phases. Before immersion, the electrodes were rinsed with deionized water and gently dried with a Kimwipe (Kimtech). The potential difference between the two electrodes immersed in the dilute phase was used as the baseline. Ag/AgCl electrodes were immersed into either the dense phase or dilute phase. Open-circuit electric potentials were recorded at room temperature as a function of time at intervals of 0.01 s for 10 s with a Metrohm Autolab PGSTAT204 station.

Alexa Fluor 647 NHS ester assay for condensate partitioning

Partitioning of negatively charged small molecules into ELP condensates was evaluated using Alexa Fluor 647 NHS ester (Thermo Fisher Scientific), a negatively charged fluorescent dye that is inert and insensitive to pH across the experimental range. The dye was added to samples at a final concentration of 200 µM and incubated for 30 min at room temperature in PhenoPlate™ 384-well microplates (Revvity) to allow equilibration. Confocal fluorescence images were acquired using a Leica Stellaris 8 Falcon equipped with an oil-immersion objective. Excitation was set to 647 nm, and emission was collected from 670-700 nm. Regions of interest (ROIs) were manually drawn around individual condensates and around adjacent dilute phase regions within the same imaging frame and focal plane (z-slice). This approach was necessary because Alexa Fluor 647 is not ratiometric and is sensitive to local concentration. The relative partition coefficient was calculated by dividing the mean pixel intensity of the condensate ROI by that of the surrounding dilute phase.

Rhodamine 6G assay for condensate partitioning

Partitioning of positively charged small molecules into ELP condensates was evaluated using Rhodamine 6G (Sigma Aldrich), a positively charged fluorescent dye that is inert and insensitive to pH across the experimental range. The dye was added to samples at a final concentration of 200 µM and incubated for 30 min at room temperature in PhenoPlate™ 384-well microplates (Revvity) to allow equilibration. Confocal fluorescence images were acquired using a Leica Stellaris 8 Falcon equipped with an oil-immersion objective. Excitation was set to 525 nm, and emission was collected from 550-570 nm. Regions of interest (ROIs) were manually drawn around individual condensates and around adjacent dilute phase regions within the same imaging frame and focal plane (z-slice). This approach was necessary because Rhodamine 6G is not ratiometric and is sensitive to local concentration. The relative partition coefficient was calculated by dividing the mean pixel intensity of the condensate ROI by that of the surrounding dilute phase.

Partition of 4-hydrazinobenzoic acid and 4-nitrobenzaldehyde in dense and dilute phase

Single-reactant samples were prepared by adding either 4-hydrazinobenzoic acid (4-HBA) or 4-nitrobenzaldehyde (4-NBA) to a solution containing ELP condensates (total volume 40 µL in PCR tubes). The solid was dissolved in DMSO and working solutions were prepared in water, yielding 10 µM 4-HBA and 40 µM 4-NBA upon mixing with 40 µL pre-equilibrated ELP condensate suspensions. $\lambda_{max}$ values of 230 nm (4-HBA) and 270 nm (4-NBA) were used for quantification.

For bulk (pre-spin) measurements, 5 µL of the condensate-containing sample was removed, mixed 1:1 with Milli-Q water (final 10 µL) to dissolve condensates, and read on NanoDrop 2000 at $\lambda_{max}$ with the instrument's 1-cm equivalent pathlength normalization enabled. Absorbance was multiplied by the 2× dilution factor. Dilute phase was separated from dense phase by centrifugation at room temperature at 20,000 × g for 30 min at 25 °C. The supernatant (dilute phase) was transferred carefully and read directly at $\lambda_{max}$. The dense-phase sequestration fraction was computed as: $f_{dense} = 1 - \frac{A_{dilute}}{A_{bulk}}$, assuming dense phase volume is negligible.

Chemical reaction between 4-hydrazinobenzoic acid and 4-nitrobenzaldehyde

Hydrazone coupling between 4-HBA and 4-NBA was used to probe partition-modulated reactivity. Solid was dissolved in DMSO and working solutions were prepared in water, yielding 10 µM 4-HBA and 40 µM 4-NBA upon mixing with 40 µL solution containing ELP condensates. Reactions were conducted at room temperature without shaking. To identify the product band, 4-HBA and 4-NBA were mixed in 1 x PBS lacking condensates and scanned from 350-600 nm on a SPARK multimode microplate reader (Tecan Life Sciences); a new maximum at 410 nm appeared after mixing, whereas no peak at 410 nm was observed prior to mixing.

For each repetition, 40 µL of condensate-containing solution was prepared and mixed with reactants. At indicated times, 5 µL of solution was mixed 1:1 with Milli-Q water to dissolve condensates and read at 410 nm against matched blanks (solution containing ELP condensates that lacks reactants and solution containing reactants and 0.25 mM ELP protein but no ELP condensates). Absorbance was measured on a NanoDrop 2000 spectrophotometer (Thermo Fisher Scientific) and multiplied by the 2× dilution factor. Initial rates were estimated from the early linear regime.

**Table S1.** Enthalpy decomposition of $\Delta\Delta G$.

| | $Cl^-$ | $Na^+$ |
|---|---|---|
| $\Delta\Delta G$ (kcal/mol) | 6.08 ± 0.08 | -4.99 ± 0.07 |
| $\Delta\Delta H$ (kcal/mol) | 8.54 ± 3.04 | -6.01 ± 1.60 |

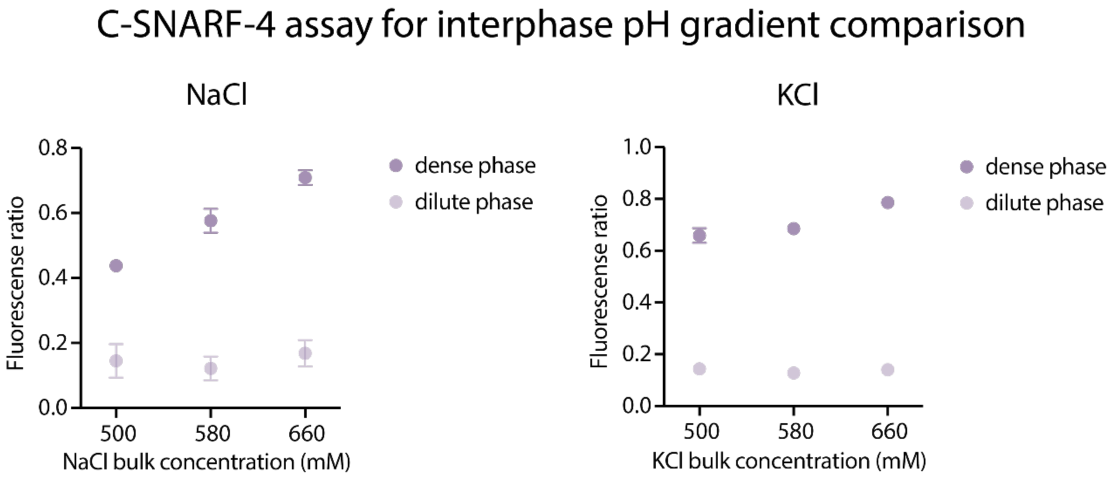


**Figure S1. Condensate salt and protein concentration collectively regulate pH capacity.** C-SNARF-4 assay for interphase pH comparison. Data represent mean ± s.d. from n = 3 independent trials.

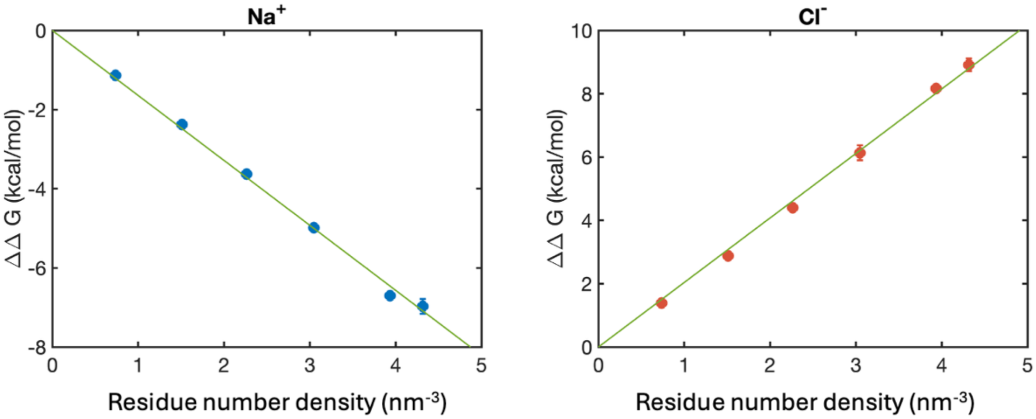


**Figure S2.** The intrinsic transfer free energy has a linear relationship with the protein volume fraction characterized by the amino acid (AA) number density.

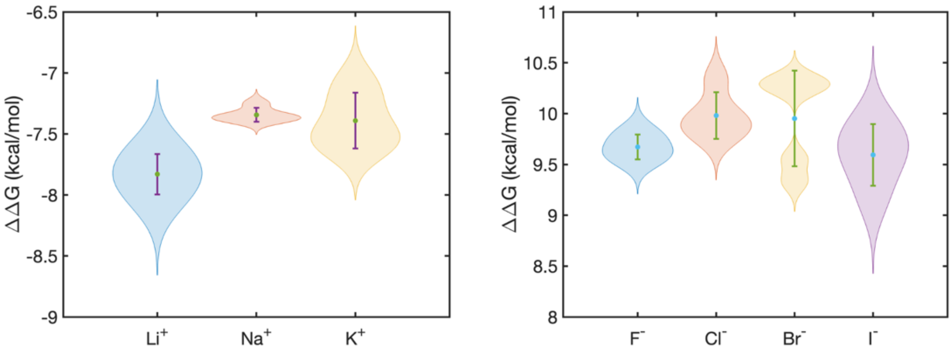


**Figure S3.** Intrinsic transfer free energies calculated from all-atom molecular dynamics simulations with amber03ws forcefields [11] with tip4p/2005 water model [12].

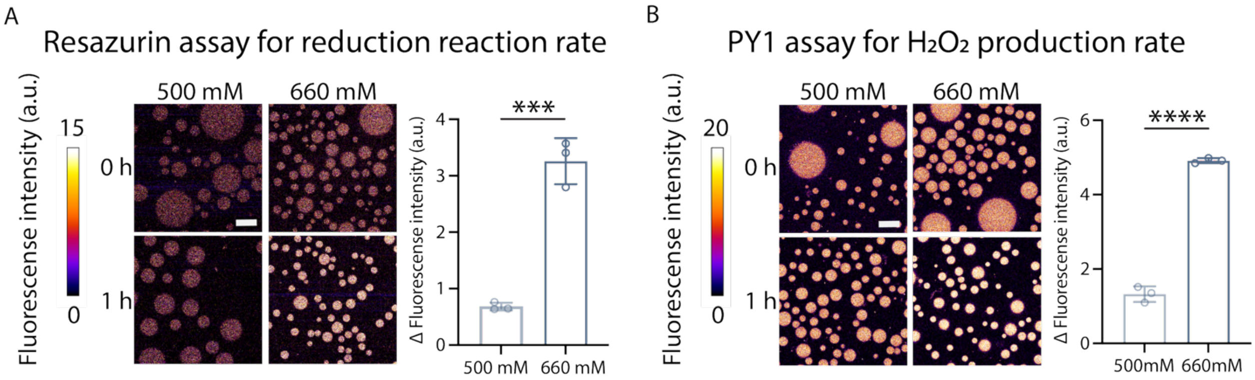


**Figure S4. ELP condensates exhibit intrinsic redox activity, modulated by salt concentration.**

**(A)** Reduction activity of condensates formed in NaCl was assessed using the fluorogenic resazurin assay. Non-fluorescent resazurin is reduced to fluorescent resorufin over time. Fluorescence within the dense phase was measured at 0 and 1 h of incubation at room temperature (no shaking), and the change in fluorescence intensity (Δ Fluorescence) was quantified to compare reduction activity across condensates formed under different conditions. Data represent mean ± s.d. from n = 3 independent samples, with >10 similar-sized condensates analyzed per sample. Scale bar, 20 μm.

**(B)** Oxidation activity of condensates formed in NaCl was measured using the Peroxy Yellow 1 (PY1) assay. In this reaction, hydroxide anions ($OH^-$) are spontaneously oxidized to hydroxyl radicals (•OH), which subsequently recombine to form hydrogen peroxide ($H_2O_2$). PY1 emits fluorescence upon reacting with $H_2O_2$, enabling quantification of oxidative capacity. Fluorescence within the dense phase was measured at 0 and 1 h of incubation at room temperature (no shaking), and the change in fluorescence intensity (Δ Fluorescence) was quantified to compare $H_2O_2$ formation rate across condensates formed under different conditions. Data represent mean ± s.d. from n = 3 independent samples, with >10 similar-sized condensates analyzed per sample. Scale bar, 20 μm.

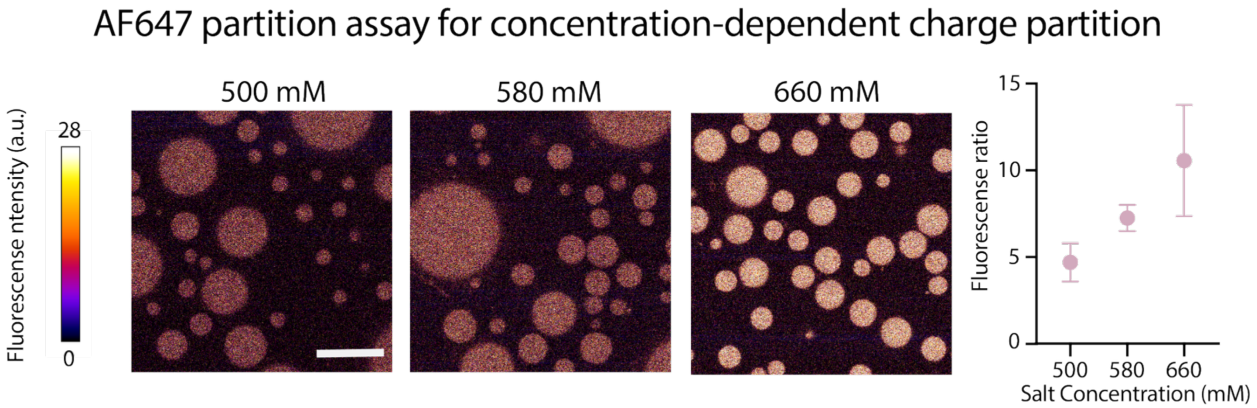


**Figure S5. ELP formed at distinct salt concentration enables different partitioning of negative charge.** Evaluation of negative charge partitioning using Alexa Fluor 647 NHS ester dye in ELP condensates formed at 500, 580, and 660 mM of NaCl. Fluorescence ratio denotes the intensity in the dense phase divided by that in the dilute phase. Data represent mean ± s.d. from $n$ = 3 independent samples; >10 condensates of similar size were analyzed per condition. Statistical analysis was performed using a two-tailed t- test: *$p < 0.05$. Scale bar, 20 µm.

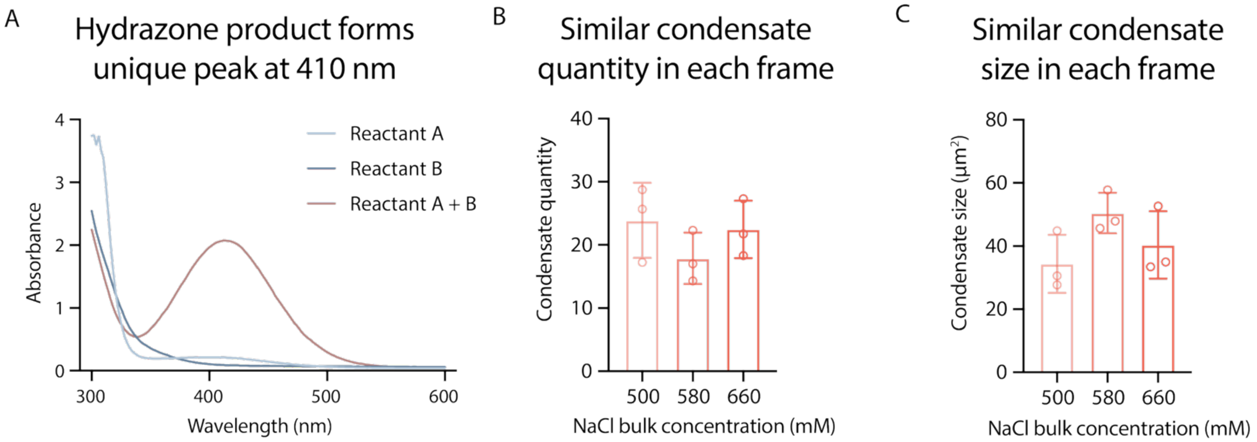


**Figure S6. Spectral confirmation and condensate uniformity for hydrazone coupling.**

**(A)** UV–Vis absorbance spectra of 1× PBS (40 μL) containing 4-hydrazinobenzoic acid (A), 4-nitrobenzaldehyde (B), or A+B immediately after mixing; $\lambda_{max}$ for A (230 nm), B (270 nm) and the hydrazone product (410 nm) are indicated.

**(B)** Condensate number per field of view (78 × 78 μm) prior to initiating reactions. Data represent mean ± s.d. from *n* = 3 independent samples; >10 condensates of similar size were analyzed per condition.

**(C)** Condensate size (projected area) per field of view (78 × 78 μm). Data represent mean ± s.d. from *n* = 3 independent samples; >10 condensates of similar size were analyzed per condition.
